# De novo single-cell biological analysis of drug resistance in human melanoma through a novel deep learning-powered approach

**DOI:** 10.1101/2025.08.18.670074

**Authors:** Sumaya Alghamdi, Turki Turki, Y-h. Taguchi

## Abstract

Elucidating drug response mechanisms in human melanoma is crucial for improving treatment outcomes. Although scRNA-seq captures gene expression at the individual cell level, existing tools to gain deeper insights into a studied population pertaining to melanoma drug resistance are far from perfect. Therefore, we propose a deep learning (DL)-based approach to unveil molecular mechanisms underlying melanoma drug resistance as follows. First, we processed two single-cell datasets related to human melanoma from GEO (GSE108383_A375 and GSE108383_451Lu) database and trained a fully connected neural network with five adapted methods (L1-Regularization, DeepLIFT, SHAP, IG, LRP) to discriminate between BRAFi-resistant and parental cell lines, followed by identifying top 100 genes. Compared to existing bioinformatics tools from a biological perspective, the presented DL-based methods identified more expressed genes in four well-established melanoma cell lines, including MALME-3M, MDA-MB435, SK-MEL-28, and SK-MEL-5. Moreover, we identified FDA-approved melanoma drugs (e.g., Vemurafenib and Dabrafenib), critical genes such as ARAF, SOX10, DCT, AXL, and key TFs including MITF and TFAP2A. From a classification perspective, we provided gene sets by all methods to three machine learning algorithms, including support vector machine, random forests, and neural networks. Results demonstrate that Integrated gradients (IG) method adapted in our DL approach contributed to 2.2% and 0.5% overall performance improvements over the best-performing baselines when using A375 and 451Lu cell line datasets.

## 1. Introduction

Understanding the mechanisms of drug response and resistance in melanoma is crucial for clinical decision making process. This knowledge contributes to narrowing down the search for candidate compounds, improving awareness and management of potential adverse reactions prior to clinical trials, and identifying possible drug targets [1]. Single-cell technologies, such as scRNA-seq, offer a powerful approach to dissect these mechanisms by providing a high-resolution view of cellular heterogeneity within tumors [2]. Research has been conducted to analyze scRNA-seq to explore essential aspects of melanoma biology, focusing particularly on drug responses and treatment resistance, aiming to reveal various molecular mechanisms. Ho et al. [3] proposed SAKE, a method specifically designed for analyzing scRNA-seq data, to tackle the analytical challenges posed by high dropout rates and complex population substructures. They employed SAKE to investigate melanoma cells resistant to BRAF inhibitors, analyzing data from the Fluidigm C1 and 10x Genomics platforms. The study discovered and recognized resistance markers, such as DCT, AXL, and NRG1, which were experimentally validated. Their analysis demonstrates that resistant cells can emerge from rare populations already present before drug application, highlighting the effectiveness of scRNA-seq in uncovering drug resistance mechanisms in melanoma. However, the study mainly focused on previously published resistance markers for BRAF inhibitors, potentially limiting new discoveries. Schmidt et al. [4] utilized scRNA-seq to investigate the heterogeneity of BRAF V600E-mutant melanoma cells under treatment with BRAF and MEK inhibitors. The study employed pseudotime trajectory and RNA velocity analyses to explore the dynamic transcriptional states associated with both treatment sensitivity and resistance.

Seven distinct cellular states were identified, with KDM5B highlighted as a pivotal marker of resistance. However, the study’s focus on a single cell line may limit the broader applicability of its findings. Zhang et al. [5] applied single-cell RNA sequencing to analyze cells from Acral Melanoma (AM) and cutaneous melanoma (CM) samples. They identified five functional cell clusters associated with key pathways, including TGF-β, Type I interferon, Wnt signaling, cell cycle, and cholesterol efflux. AM showed a more immunosuppressive microenvironment, with reduced cytotoxic CD8+ T cells and elevated expression of exhaustion markers PD-1 and TIM-3. While the study provides valuable insights into immune heterogeneity, its limited patient sample size and subtype-specific focus may constrain broader applicability.

Egan et al. [6] developed a regulon-based approach to analyze immune cell states in melanoma utilizing scRNA-seq data. The researchers employed clustering, and differential expression analysis with other computational techniques to identify four predominant immune cell states: exhausted T cells, monocyte lineage cells, memory T cells, and B cells. These identified states were subsequently validated using bulk RNA-seq data and were found to be associated with differential responses to immune checkpoint inhibitors (ICIs). The study further elucidated an interaction between monocyte lineage cells and T cell exhaustion, suggesting that monocyte lineage cells may drive T cells into a state of terminal exhaustion through mechanisms involving antigen presentation and chronic inflammation pathways. This regulon-based analysis offers a robust approach for predicting responses to ICIs; however, it is important to note the limitations associated with the small sample size of 19 and potential information loss resulting from data reduction. Li et al. [7] characterized the immune landscape of AM, a rare subtype of melanoma, utilizing scRNA-seq data. The analysis encompassed clustering, differential expression analysis, and cell-cell interaction analysis, followed by a comparative examination with non-acral melanoma. The study revealed a suppressed immune environment in AM, characterized by a reduced presence of immune cells, including CD8+ T cells and natural killer (NK) cells, in contrast to non-acral melanoma. Furthermore, the research stated several immune checkpoints, such as PD-1, LAG-3, VISTA, and TIGIT, as potential targets for immunotherapy. However, despite providing valuable insights, the limited sample size and the primary focus on the comparison between AM and non-acral melanoma constrain the generalizability of the findings. Zhang et al. [8] aimed to investigate CD3ζ as a predictive biomarker for resistance to PD-1 inhibitors in melanoma. The researchers conducted an analysis of scRNA-seq data to identify CD3ζ as a key marker. Low expression levels of CD3ζ demonstrated a strong correlation with poor prognosis, diminished immune cell infiltration, and resistance to PD-1 inhibitors. In melanoma, the augmentation of CD3ζ expression was associated with enhanced efficacy of the PD-1 inhibitor nivolumab, indicating that CD3ζ may serve as a therapeutic target to counteract resistance. However, the study’s narrow focus on CD3ζ may have overlooked other pertinent factors contributing to resistance.

Other studies have been conducted an analysis using both scRNA-seq and bulk RNA-seq to investigate drug response in melanoma. Bakr et al. [9] investigated the role of nicotinic acetylcholine receptors (CHRNs) in melanoma metastasis by first utilizing publicly available bulk RNA-seq data to conduct differential expression, correlation, and survival analyses. The results indicated that CHRNA1 is significantly expressed in metastatic melanoma and is associated with metastasis via pathways such as ZEB1 and Rho/ROCK. Further analyses demonstrated that CHRNA1 is connected to key prognosis-related genes, including DES, FLNC, CDK1, and CDC20. These findings were then validated using scRNA-seq data, confirming CHRNA1’s role as a potential prognostic marker for metastatic melanoma. However, the focus on CHRNA1 might oversimplify the complex nature of metastasis. Wang et al. [10] examined the role of Aurora kinase B (AURKB) in suppressing the immune response against melanoma by initially conducting differential expression, immune infiltration, and survival analyses using bulk RNA-seq data. Their findings suggested that AURKB inhibits immune activity by modulating signaling pathways between tumor cells and lymphocytes. The researchers further validated these results using scRNA-seq data, confirming that inhibition of AURKB through Tozasertib reduced tumor growth by decreasing regulatory T cells and activating CD8+ T cells. Despite these promising results, variability in patient responses to Tozasertib highlights the need for further research to optimize its therapeutic application

In addition to traditional bioinformatics tools, recent studies have applied artificial intelligence (AI), including machine learning, to enhance the analysis of scRNA-seq in melanoma. Chen et al. [11] identified three key prognostic genes (SLC25A38, EDNRB, LURAP1) associated with Uveal Melanoma (UM) metastasis by applying machine learning to single-cell and bulk RNA-seq data. Prognostic genes were identified using the log-rank test and univariate Cox regression, and prognostic models were developed with LASSO and multivariate Cox regression analyses. Single-cell analysis revealed that high-risk patients had increased infiltration of CD8+ T cells, which were largely exhausted and dysfunctional. The study also highlighted altered cell– cell communication, especially through CD99 and MHC-I pathways, as potential contributors to metastasis. Pinhasi and Yizhak developed a machine learning approach, PRECISE, to predict immunotherapy response in melanoma using single-cell RNA-seq data from tumor-infiltrating immune cells. Their approach combined XGBoost with Boruta feature selection and SHAP analysis to identify an 11-gene signature (including GAPDH, STAT1, CD38) predictive of immune checkpoint inhibitor response. They also used reinforcement learning to quantify each cell’s contribution to prediction, achieving an AUC of 0.89 and validating their model across multiple cancer datasets. Table1 provides a summary of the related works included in this study.

**Table 1:** Summary of the related works pertaining to melanoma when compared against our proposed work.

| Ref. | Year | Objective | Data Source | Main Findings |
| --- | --- | --- | --- | --- |
| [3] | 2018 | Identifying resistance markers to BRAF inhibitors in melanoma. | scRNA-seq | Known markers (DCT, AXL, NRG1). |
| [4] | 2022 | Analyzing resistance in BRAF V600E melanoma cells. | scRNA-seq | KDM5B as marker; distinct sensitivity/resistance states. |
| [5] | 2022 | Investigate tumor heterogeneity and immune environment in AM and CM. | scRNA-seq | Five cell clusters (e.g., TGF-β, cell cycle) associated with good melanoma prognosis. |
| [6] | 2022 | Characterizing immune cell states and predicting responses to ICIs, | scRNA-seq | Four immune states; patient clustering into response groups. |
| [7] | 2022 | Characterizing the cellular and immune landscape in AM. | scRNA-seq | AM has fewer immune cells; immune checkpoints (PD-1) as therapeutic targets. |
| [8] | 2023 | Identifying biomarkers predicting resistance to PD-1 inhibitors. | scRNA-seq | CD3ζ predicts PD-1 resistance. |
| [9] | 2023 | Investigating CHRNA1's role in melanoma metastasis. | scRNA-seq/<br>bulk | CHRNA1 is highly expressed in metastasis. |
| [10] | 2024 | Assessing the impact of AURKB inhibition on melanoma immune response. | scRNA-seq/<br>bulk | AURKB inhibition enhances anti-tumor immunity;Tozasertib reduces tumor growth. |
| [11] | 2024 | Identifying prognostic genes for UM metastasis using ML. | scRNA-seq/<br>bulk | Three prognostic genes identified (SLC25A38, EDNRB, LURAP1). |
| [12] | 2025 | Enhancing the prediction of treatment outcomes in melanoma using ML. | scRNA-seq | 11-gene signature (e.g., GAPDH, STAT1); |
| Proposed | 2025 | Unveiling various molecular mechanisms underlying melanoma drug resistance; performance validation from classification and biological perspectives against baseline methods | scRNA-seq | -Adapting five DL-based methods: L1-Regularization, DeepLIFT, SHAP, IG, LRP.<br>-Biological Perspective: More expressed genes, Approved FDA drugs (Vemurafenib and Dabrafenib), TFs as MITF and TFAP2A<br>-Classification Results: 2.2% and 0.5% performance improvements over baseline methods |

While bioinformatics approaches are currently central to understanding melanoma drug mechanisms, they often rely on existing tools and do not exploit the potential of artificial intelligence. Furthermore, existing AI-driven approaches have yet to fully explore the potential of combining deep learning (DL) with single-cell analysis and thereby are far from perfect. This study presents a novel AI-driven computational approach that integrates adapted DL-based methods followed by enrichment analysis, specifically designed to enhance the analysis of single-cell data to uncover deeper biological insights. Our approach aims to facilitate the practical application of AI in clinical settings and enable the proactive identification of therapeutic targets associated with effective treatment responses in melanoma. The key innovations of this work include:

1. DL-powered computational approach: We introduce a DL-based approach for gene selection with enrichment analysis to analyze single-cell gene expression data, unveiling mechanism underlying melanoma drug response, critical genes, drugs, therapeutic targets, and other critical factors such as key pathways and regulators associated with melanoma treatment outcomes.
2. Model training: We downloaded and processed single-cell gene expression datasets related to melanoma treatment response from the GEO under accession numbers GSE108383_A375 and GSE108383_451Lu, referred to as A375 cell lines and 451Lu cell lines, respectively. Then, a fully connected deep neural network was trained to learn complex non-linear relationships in the single-cell gene expression data, automatically engineer informative genes, and implicitly reduce dimensionality, enabling more accurate identification of crucial genes for predicting drug response.
3. Gene ranking and selection: Adapting DL-based gene selection methods—including L1-Regularization, DeepLIFT, SHAP, Integrated Gradients (IG), and Layer-wise Relevance Propagation (LRP)—to rank genes based on their importance to the model predictions. Subsequently, we selected the top-ranked genes determined by each method for further enrichment analysis, including using Enrichr and Metascape to uncover the underlying biological pathways and processes associated with drug response and resistance in melanoma.
4. Results from biological perspective: Our developed DL-based approach demonstrated superior performance when compared to baseline methods across the two datasets. Notably, DeepLIFT and IG produced the highest performance on the first dataset, whereas both DeepLIFT and IG were most effective for the second, with higher number of expressed genes in established cell lines (i.e., MALME-3M, MDA-MB435, SKMEL28, and SKMEL5). Additionally, our methods successfully identified several drugs, including FDA-approved melanoma treatments like Vemurafenib and Dabrafenib, significant genes (e.g., ARAF, SOX10, SLC7A5, DCT, and AXL), and TFs (such as MITF, RELA, E2F1, and TFAP2A).
5. Results from classification perspective: Providing the gene sets delivered by our DL-based methods and existing baseline methods to three machine learning algorithms, evaluated using the average balanced accuracy (BAC) and F1. Compared to the best-performing baseline (LIMMA) in the A375 cell line dataset, IG coupled with machine learning algorithms had performance improvements of 2.2% and 2.0%, measured using F1 and BAC, respectively. For the 451Lu cell line dataset, IG coupled with machine learning algorithms resulted in 0.5% and 0.3% performance improvements when employing F1 and BAC performance measures, respectively, over the best-performing baseline (t-test).

## 2. Materials and Methods

### 2.1. Single-cell Gene Expression Profiles

In this study, we utilized two scRNA-seq datasets, GSE108383_A375 and GSE108383_451Lu, obtained from the gene expression omnibus (GEO) under accession number GSE108383. These datasets are derived from the A375 and 451Lu melanoma cell lines, which both carry the BRAF V600E mutation, a prevalent driver mutation in melanoma [13]. We refer to GSE108383_A375 and GSE108383_451Lu as A375 cell line and 451Lu cell line, respectively. Analysis of these datasets in the presence of sophisticated computational methods enable us to unveil various mechanisms leading to drug resistance in melanoma patients by providing a detailed view of transcriptional heterogeneity at the single-cell level.

As mentioned, all datasets consist of single-cell gene expression data, where each sample *x*_*i*_ *ϵ* R^*n*^ composed of the expression of *n* genes, and *y*_*i*_ *ϵ* {0,1} is the binary label indicating the cell state (0 for parental and 1 for resistant). Table 2 provides an overview of these two datasets.

**Table 2:**
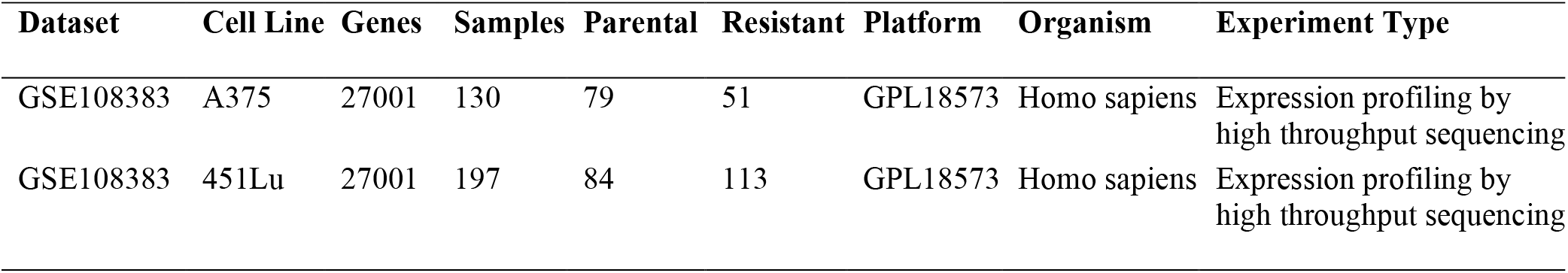
Overview of scRNA-seq datasets downloaded from the gene expression omnibus (GEO) database.

| Dataset | Cell Line | Genes | Samples | Parental | Resistant | Platform | Organism | Experiment Type |
| --- | --- | --- | --- | --- | --- | --- | --- | --- |
| GSE108383 | A375 | 27001 | 130 | 79 | 51 | GPL18573 | Homo sapiens | Expression profiling by high throughput sequencing |
| GSE108383 | 451Lu | 27001 | 197 | 84 | 113 | GPL18573 | Homo sapiens | Expression profiling by high throughput sequencing |

### 2.2. Computational Approach

The proposed approach involves data preparation and gene selection using DL-based methods, including L1 Regularization, DeepLIFT, SHAP, IG and LRP. These methods were adapted to single-cell gene expression data to identify the most important genes that distinguish between parental and resistant cell states. The selected top genes were then subjected to enrichment analysis using Enrichr and Metascape to uncover the underlying biological pathways and processes associated with drug response and resistance in melanoma (Figure 1).

**Figure 1:**
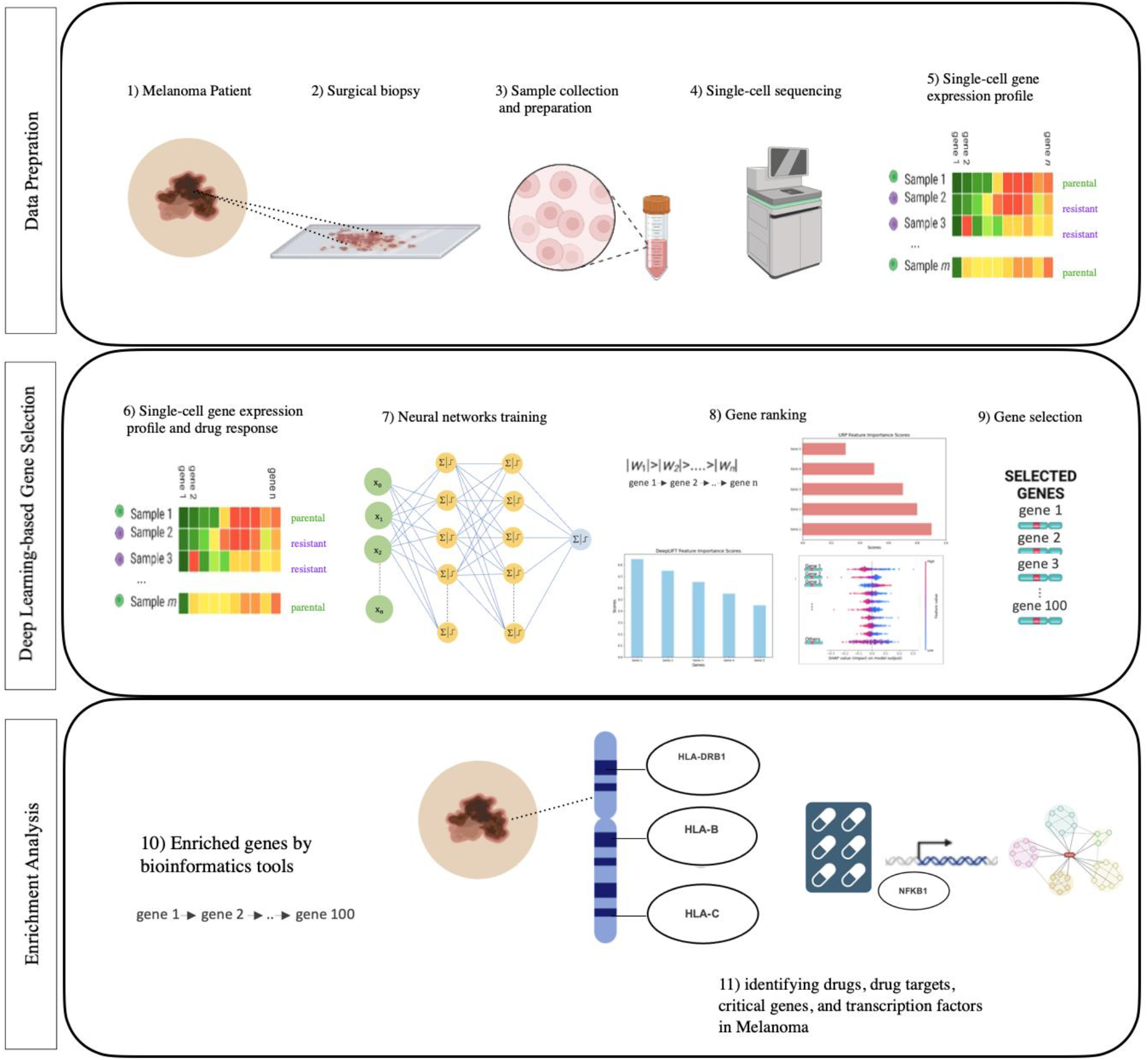
Flowchart of the DL-powered computational approach identifying drugs, drug targets, critical genes, and transcription factors in melanoma.

#### 2.2.1. Data Preparation

As part of the data preparing step, biopsy samples were obtained from melanoma patients. Then, collected samples were prepared and provided to a single-cell technology, measuring the gene expression levels at single-cell resolution. The data were then annotated according to their respective cell states with labels assigned as 0 for parental and 1 for resistant [14].

#### 2.2.2. Deep Learning-based Gene Selection

##### Neural Network Model

A fully connected feedforward neural network [15] was trained on the single-cell data. Before the training, we performed min-max scaling to normalize the expression levels of each gene to the range [0,1], ensuring that all genes are comparable. Following that, the datasets were split into training and testing sets, ensuring a balanced representation of both cell states in each subset. The data was then fed to the fully connected feedforward neural network which consists of two hidden layers with ReLU activation function [16] and dropout layer [17] to prevent overfitting. The first hidden layer transforms the input data using a weight matrix w_1_ *ϵ R*^*p*×*n*^ and a bias vector *b*_*1*_ *ϵ R*^*p*^, where *n* is the number of input genes and *p* is the number of neurons in the hidden layer (*p* =128). This transformation is followed by a ReLU activation function and a dropout layer with a rate of 0.5. The second hidden layer applies a similar transformation, and the final output layer uses a sigmoid activation function to predict the probability that a given sample belongs to one of the two classes (see Figure 2).

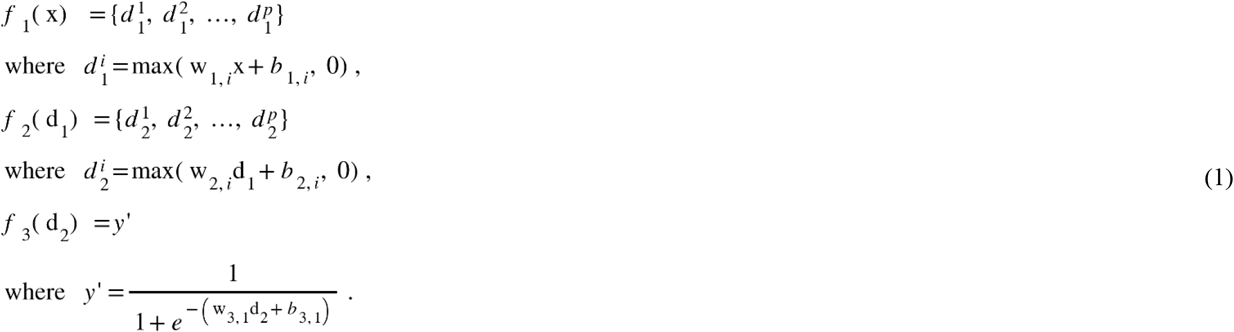

**Figure 2:**
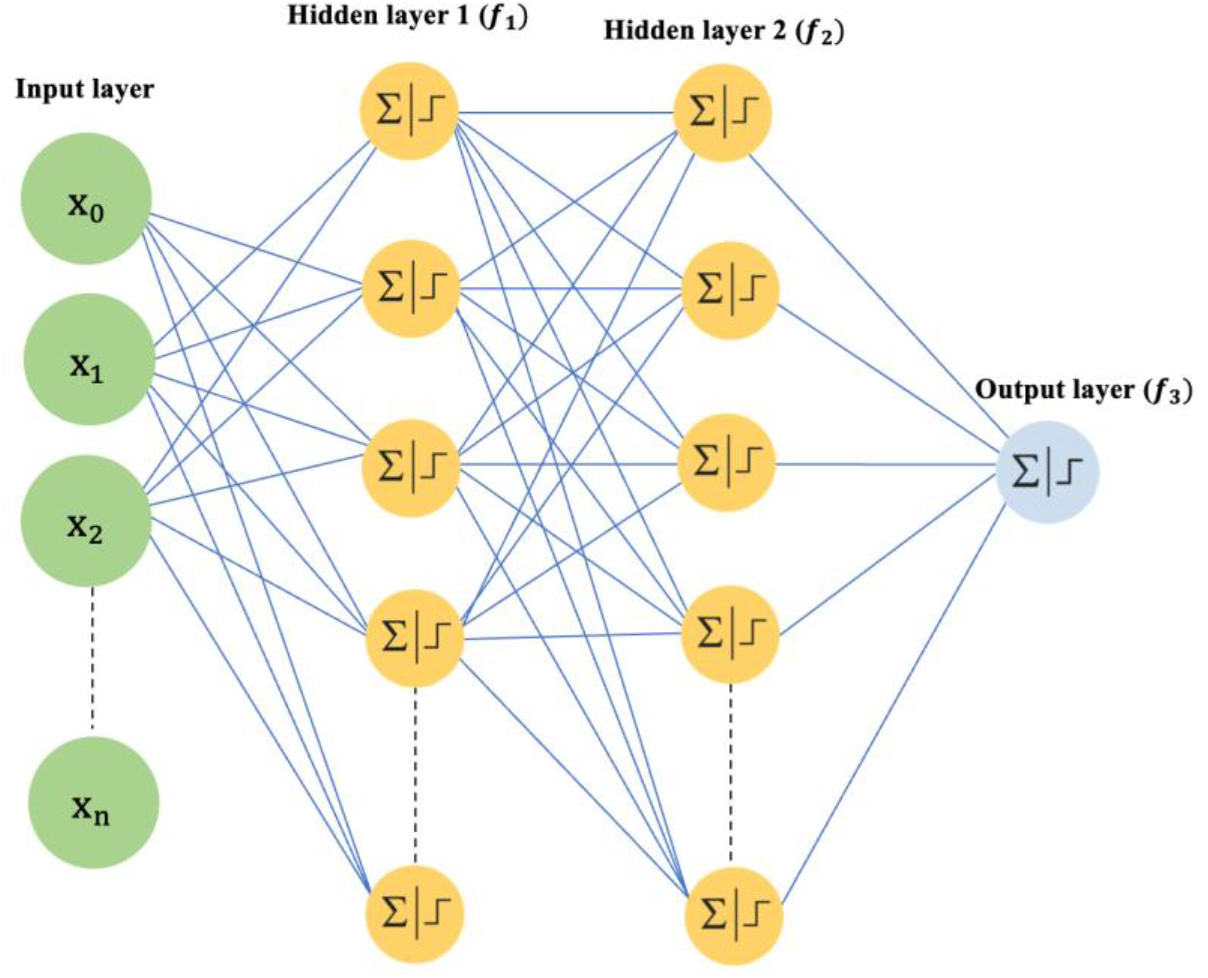
Architecture of the fully connected feedforward neural network in the proposed approach.

The overall model is defined as a composition of these transformations as:

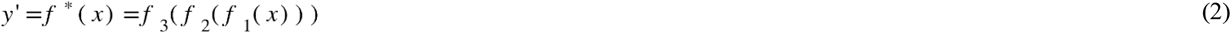

The model was trained using binary cross-entropy (BCE) loss, defined as:

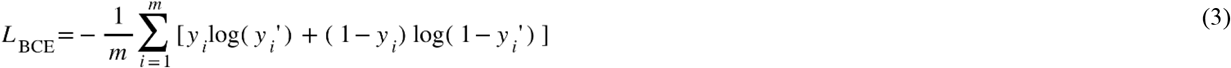

##### L1-Regularization for Gene Selection

L1-regularization was used during training to encourage sparsity in the learned weights and prevent overfitting. This technique adds a penalty term to the loss function that reflects the magnitude of the model’s weights. This encourages the model to learn a sparse representation, effectively driving the weights of less informative genes toward zero [15, 18]. For gene selection, we focused on the weights of the first hidden layer, which directly connects the input genes to the neurons of that layer. The L1-regularization term is defined as:

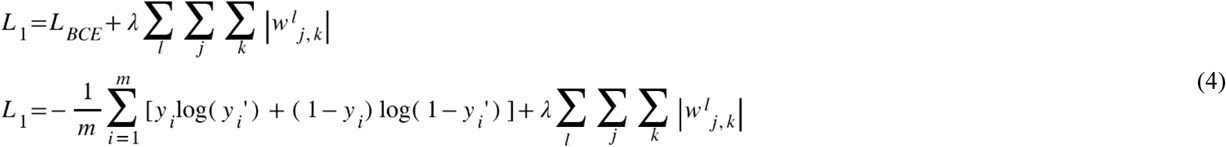

where *λ* is the regularization strength, 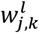 is the weight within the weight matrix in the *l*th hidden layer. We ranked the genes by summation of the absolute values of their corresponding weights across all neurons in the first layer and then selected the top 100 genes.

###### Deep Learning Important Features (DeepLIFT)

DeepLIFT is a method that explains predictions of complex deep neural networks by decomposing them into contributions of individual neurons through back-propagation. For a particular prediction, it generates local explanations by calculating the difference between the actual output and a reference output, relative to the differences between actual and reference inputs [19]. Formally, let *o* represent some target output neuron of interest and let *x*_1_, *x*_2_, …, *x*_*n*_ represent some neurons in some hidden layer. Let *f* (*x*) and *f* (*x′*) correspond to the activation of a particular neuron and the reference activation, respectively. DeepLIFT calculates contributions scores satisfying the summation-to-delta property as follows:

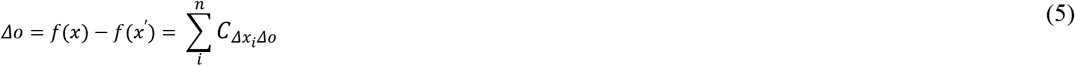

where 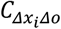 denotes the contribution score in each neuron *x*_*i*_. Let *o* and *o*^0^ correspond to target output and its reference activation, respectively. Then, the difference-from-reference is computed as *Δo* = *o* − *o*^0^[19].

To quantify this contribution, DeepLIFT defines a multiplier as follows:

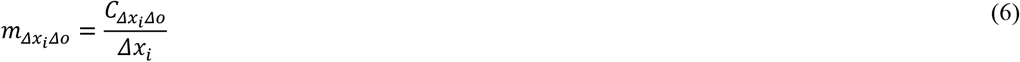

These multipliers are propagated recursively through the network layers by applying a chain rule:

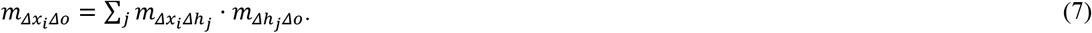

where *h*_*j*_ represents neurons in intermediate (hidden) layers. After computing the contributions for each input across all samples using DeepLIFT, we summed their absolute values to obtain a global importance score. We then ranked the genes by these scores and selected the top 100 genes for further analysis.

##### SHapley Additive exPlanations (SHAP)

SHAP is a method derived from game theory that is used to explain the output of machine learning models. It is built on the concept of the shapley value, which is a solution in coalitional game theory used to fairly distribute payouts among players in a coalition. SHAP adapts this idea to machine learning by attributing the contribution of each gene to a model’s prediction [20]. The SHAP value *ϕ*_*j*_ for a given gene *j* in a prediction *q* can be defined mathematically as:

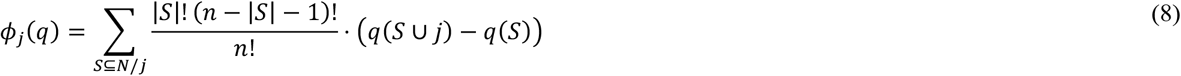

where *n* is the total number of genes, *S* ⊆ *N*/*j* represents all possible subsets of genes excluding *j*, (*q*(*S* ∪ *j* − *q*(*s*)) is the marginal contribution of gene *j*, i.e., the difference in predictions when the gene *j* included versus excluded, and 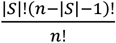 is the weighting factor for the marginal contributions, ensuring fairness by considering all possible gene subsets.

After calculating the SHAP value for each gene, genes were ranked based on the sum of their absolute SHAP values across all samples, and the top 100 were selected as the most influential genes obtained from the model.

##### Integrated Gradients (IG)

IG is a gradient-based explainability method that helps uncover the relative importance of input genes in predictions by a model. It compares the output of a model for a given input to its output for a baseline value. This baseline represents a neutral state, serving as a starting point for the explanation [21]. The method calculates gene contributions by integrating the gradients of the model’s output along a straight path from the baseline input to the actual input. Mathematically, this is represented as:

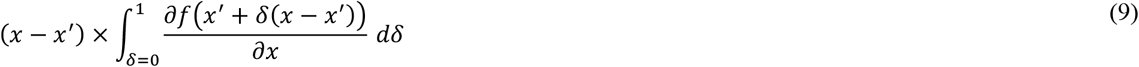

where *f* denotes the trained neural network model, *x* is the input gene, *x*^*′*^ is the baseline, *δ* is a scalar interpolating between them, and the integral accumulates the gradients along the path.

We computed the absolute IG scores for each gene and summed them across all samples. The top 100 genes were then selected as the key contributors to the model’s predictions.

##### Layer-wise Relevance Propagation (LRP)

LRP is an explainability method designed to trace the contribution of input genes to final prediction of the model by redistributing the output relevance backward through the network layers. This redistribution follows predefined propagation rules that ensure the total relevance remains conserved across layers, matching the output of the model at the final layer [22]. One of the fundamental rules in LRP is the simple rule (LRP-0), which redistributes the relevance 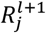 from layer *l* + 1 to layer *l* proportionally, based on each contribution of the input. The rule is defined as:

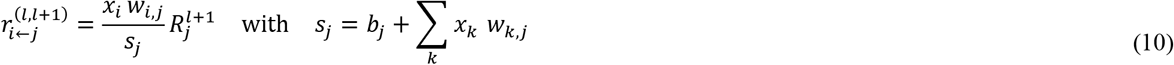

here, 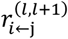 denotes the relevance passed from neuron *j* in layer *l* + 1 to neuron *i* in layer *l*; *x*_*i*_ is the activation of neuron *i*; *w*_*i,j*_ is the weight connecting neuron *i* in layer *l* to neuron *j* in layer *l* + 1; *s*_*j*_ is the total pre-activation of neuron *j*, including its bias term *b*_*j*_ and the weighted sum of all inputs to *j*, and 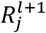 is the relevance of neuron *j* in layer *l* + 1. After computing the relevance scores for each input gene, we summed their absolute values across all samples. The top 100 genes were then selected as the most significant contributors to the model’s predictions.

#### 2.2.3. Enrichment Analysis

For the enrichment analysis, we uploaded the top significant genes in A375 cell line and 451Lu cell line as input to Enrichr and Metascape [23,24]. These genes were ranked based on their importance as determined by each method. Subsequently, we analyzed the results to interpret and identify biologically meaningful terms, such as key expressed genes, drugs, therapeutic targets, transcription factors, and other relevant categories. The findings from this analysis are presented in the next section.

## 3. Experiments and Results

### 3.1. Experimental Methodology

We compared our DL-based approach against the following baseline methods (LIMMA [25], SAM and t-test [26,27]. The input to these studied methods is our labeled gene expression of both datasets. For LIMMA, SAM, and t-test baseline methods, genes were selected based on significantly adjusted p-values < 0.01.

To perform enrichment analysis and evaluate the results from a biological perspective, we uploaded genes obtained from each method to Enrichr (https://maayanlab.cloud/Enrichr/, accessed on 3 February 2025) and Metascape (https://metascape.org/gp/index. html, accessed on 15 March 2025). When retrieved terms are related to melanoma, a method that has terms associated with more genes is considered the superior method. Furthermore, we assessed the performance from the classification perspective, reporting balanced accuracy (BAC) and F1 as our performance measures.

In this study, we utilized Python to run the experiments of our DL-based gene selection methods [28]. Specifically, for our DL-based approach, we utilized PyTorch, a widely used library for developing and training neural networks [29]. To identify important genes, DeepLIFT, IG, and LRP were implemented using the Captum library [30]. Additionally, SHAP was separately installed and imported as a standalone Python package, and SHAP’s Gradient Explainer was specifically utilized for interpreting the model [31]. Moreover, we incorporated L1-Regularization, implemented directly within PyTorch, to encourage sparsity in the model’s weights and facilitate gene selection by identifying the most important input genes.

For the bioinformatics tools, we employed the LIMMA package in R [32], utilizing the lmFit and eBayes functions [25] for differential expression analysis, and the siggenes package [33] to run SAM. In terms of the t-test, we employed the t-test function within the stats package [27]. For LIMMA, SAM, and t-test, to compute adjusted p-values, we employed the p.adjust function with the BH correction.

### 3.2. Biological Results

#### 3.2.1 A375 Cell Line

Table 3 shows the number of expressed genes in the retrieved terms (i.e., melanoma cell lines) from Enrichr within the NCI-60 cancer cell lines category. A higher number of expressed genes indicates superior performance of the computational method. Among all methods of our DL-based approach and the existing approaches, DeepLIFT performed better than others, obtaining a total of 34 expressed genes within the retrieved melanoma cell lines. Specifically, seven genes (ARPC1B; HEXA; HMG20B; SOX10; ATP6V0C; LY6E; GAS7) were expressed within MALME-3M, fourteen genes (SLC35A2; PYGB; ACOT7; GPAA1; TWF2; IMP4; SOX10; RRP7A; SEC61A1; BIN3; SLCO4A1; ST3GAL4; HMG20B; RALY) were expressed within MDA-MB435, seven genes (SLC35A2; MIA; HEXA; ST3GAL4; HMG20B; SOX10; HLA-DQB1) were expressed within SKMEL28, and six genes (PYGB; BIN3; ENTPD6; HMG20B; SOX10; LY6E) were expressed within SKMEL5. The second-best method is IG, obtaining a total of 32 expressed genes within the retrieved melanoma cell lines. In terms of MALME-3M, seven genes (ARPC1B; HEXA; HMG20B; SOX10; ATP6V0C; LY6E; GAS7) were expressed, fourteen genes (SLC35A2; PYGB; ACOT7; GPAA1; GPS1; PRKCD; IMP4; SOX10; RRP7A; SEC61A1; BIN3; ST3GAL4; HMG20B; RALY) were expressed within MDA-MB435, and five genes (SLC35A2; MIA; ST3GAL4; HMG20B; SOX10) were expressed in SKMEL28, and six genes (SDCBP; PYGB; BIN3; HMG20B; SOX10; LY6E) were expressed within SKMEL5. Both SAM and the t-test identified 30 expressed genes, SHAP, LRP, and LIMMA each identified 29 expressed genes, whereas L1-Regularization identified 22 expressed genes. In Supplementary DataSheet1_A, we include all genes produced via each computational method, provided to Enrichr and Metascape. Moreover, Supplementary Table1_A lists all enrichment analysis results for NCI-60 Cancer Cell Lines, obtained from Enrichr.

**Table 3:** NCI-60 Cancer Cell Lines via Enrichr according to the genes produced by each method when using A375 Cell Line.

| Method | Rank | Term | Overlap | P-value | Adjusted P-value |
| --- | --- | --- | --- | --- | --- |
| **L1-Regularization** | 6 | MALME-3M | 5/366 | $3.67 \times 10^{-2}$ | $4.90 \times 10^{-1}$ |
| | 3 | MDA-MB435 | 8/621 | $1.26 \times 10^{-2}$ | $3.37 \times 10^1$ |
| | 31 | SKMEL28 | 3/436 | $3.73 \times 10^{-1}$ | $8.96 \times 10^{-1}$ |
| | 2 | SKMEL5 | 6/354 | $8.73 \times 10^{-3}$ | $3.37 \times 10^{-1}$ |
| **DeepLIFT** | 4 | MALME-3M | 7/366 | $2.42 \times 10^{-3}$ | $3.68 \times 10^{-2}$ |
| | 1 | MDA-MB435 | 14/621 | $2.53 \times 10^{-6}$ | $1.67 \times 10^{-4}$ |
| | 7 | SKMEL28 | 7/436 | $6.28 \times 10^{-3}$ | $5.92 \times 10^{-2}$ |
| | 9 | SKMEL5 | 6/354 | $8.73 \times 10^{-3}$ | $6.40 \times 10^{-2}$ |
| **SHAP** | 5 | MALME-3M | 6/366 | $1.02 \times 10^{-2}$ | $1.55 \times 10^{-1}$ |
| | 1 | MDA-MB435 | 11/621 | $2.74 \times 10^{-4}$ | $2.09 \times 10^{-2}$ |
| | 7 | SKMEL28 | 6/436 | $2.22 \times 10^{-2}$ | $2.00 \times 10^{-1}$ |
| | 4 | SKMEL5 | 6/354 | $8.73 \times 10^{-3}$ | $1.55 \times 10^{-1}$ |
| **IG** | 4 | MALME-3M | 7/366 | $2.42 \times 10^{-3}$ | $3.91 \times 10^{-2}$ |
| | 1 | MDA-MB435 | 14/621 | $2.53 \times 10^{-6}$ | $1.77 \times 10^{-4}$ |
| | 9 | SKMEL28 | 5/436 | $6.77 \times 10^{-2}$ | $4.32 \times 10^{-1}$ |
| | 6 | SKMEL5 | 6/354 | $8.73 \times 10^{-3}$ | $1.02 \times 10^{-1}$ |
| **LRP** | 3 | MALME-3M | 7/366 | $2.42 \times 10^{-3}$ | $5.25 \times 10^{-2}$ |
| | 1 | MDA-MB435 | 13/621 | $1.31 \times 10^{-5}$ | $8.54 \times 10^{-4}$ |
| | 13 | SKMEL28 | 4/436 | $1.75 \times 10^{-1}$ | $7.90 \times 10^{-1}$ |
| | 6 | SKMEL5 | 5/354 | $3.25 \times 10^{-2}$ | $3.52 \times 10^{-1}$ |
| **LIMMA** | 7 | MALME-3M | 6/366 | $1.02 \times 10^{-2}$ | $9.17 \times 10^{-2}$ |
| | 2 | MDA-MB435 | 12/621 | $6.28 \times 10^{-5}$ | $1.98 \times 10^{-3}$ |
| | 16 | SKMEL28 | 4/436 | $1.75 \times 10^{-1}$ | $6.87 \times 10^{-1}$ |
| | 4 | SKMEL5 | 7/354 | $2.00 \times 10^{-3}$ | $3.16 \times 10^{-2}$ |
| **SAM** | 8 | MALME-3M | 6/366 | $1.02 \times 10^{-2}$ | $7.00 \times 10^{-2}$ |
| | 1 | MDA-MB435 | 13/621 | $1.31 \times 10^{-5}$ | $7.22 \times 10^{-4}$ |
| | 14 | SKMEL28 | 4/436 | $1.75 \times 10^{-1}$ | $6.86 \times 10^{-1}$ |
| | 5 | SKMEL5 | 7/354 | $2.00 \times 10^{-3}$ | $2.21 \times 10^{-2}$ |
| **t-test** | 8 | MALME-3M | 6/366 | $1.02 \times 10^{-2}$ | $7.13 \times 10^{-2}$ |
| | 1 | MDA-MB435 | 13/621 | $1.31 \times 10^{-5}$ | $7.36 \times 10^{-4}$ |
| | 14 | SKMEL28 | 4/436 | $1.75 \times 10^{-1}$ | $6.98 \times 10^{-1}$ |
| | 4 | SKMEL5 | 7/354 | $2.01 \times 10^{-3}$ | $2.81 \times 10^{-2}$ |

Figure 3a presents a visualization pertaining to the intersected gene sets by each computational methods within our approach as well as traditional bioinformatics tools. A total number of 19 and 10 unique genes were attributed to DeepLIFT and IG, respectively. In contrast, traditional statistical methods such as SAM, and t-test demonstrated more conservative behavior, both identifying only two unique genes, while LIMMA identified no unique genes. It can also be seen that the number of common genes is upper bounded by 34 (intersection of t-test, SAM, and LIMMA) and lower bounded by 1. These results demonstrate that our computational methods produced different gene sets when compared to existing tools. In Supplementary DataSheet1_B, we provide the genes based on the UpSet plot showing in Figure 3a.

**Figure 3:**
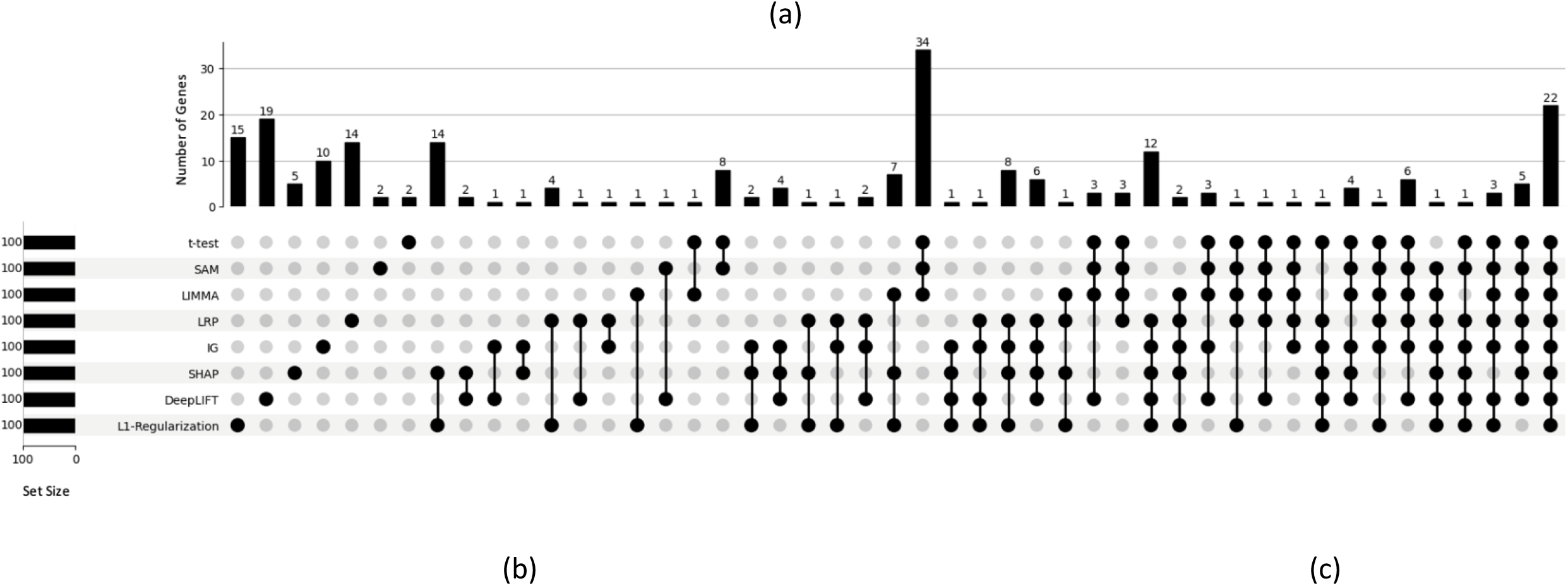
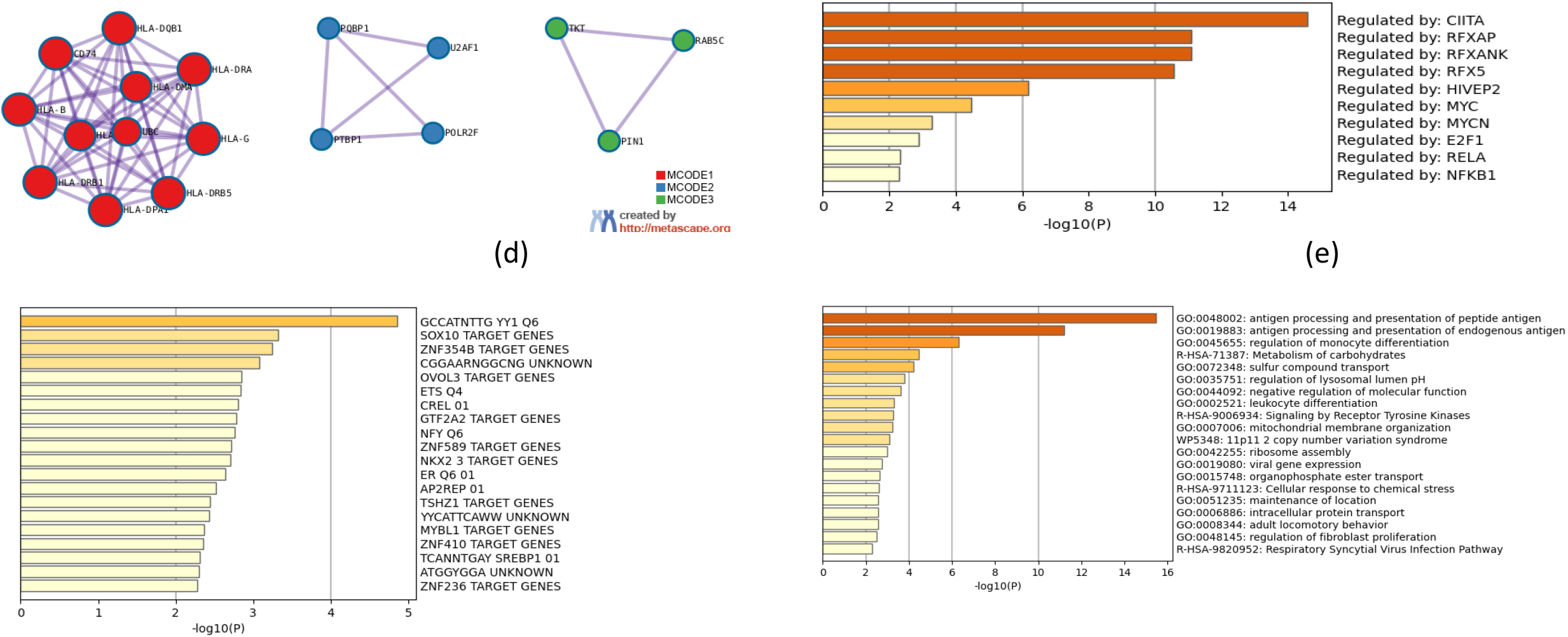
(a) UpSet plot of gene lists provided by all the computational methods when using A375 Cell Line. (b) Three clusters identified in PPI network pertaining to A375 Cell Line. (c) Ten transcription factors, according to Metascape, obtained when coupled with genes from DeepLIFT. (d) Enrichment analysis pertaining to twenty transcription factor targets. (e) Process and pathway enrichment analysis provided by Metascape according to produced genes via DeepLIFT when using A375 Cell Line.

Since DeepLIFT outperformed all the other methods, we provided the top 100 genes identified by DeepLIFT to Metascape for additional enrichment analysis. Figure 3b shows the three clusters identified in the PPI. The following 18 genes were obtained from the PPI network: HLA-B, CD74, HLA-G, HLA-DQB1, HLA-DMA, HLA-DRB5, HLA-C, HLA-DRA, UBC, HLA-DPA1, HLA-DRB1, PQBP1, PTBP1, POLR2F, U2AF1, RAB5C, TKT, and PIN1. Figure 3c reports 10 transcription factors (TFs): CIITA, RFXANK, RFXAP, RFX5, HIVEP2, MYC, MYCN, E2F1, RELA, NFKB1. CIITA plays a role as a transcriptional coactivator in melanoma, particularly by regulating immune-related genes (like MHC II) and influencing tumor immunogenicity [34]. MYC has been reported as a key oncogenic transcription factor in melanoma that promotes tumor growth and therapy resistance [35,36]. E2F1 functions as a transcription factor that regulates genes involved in melanoma cell proliferation, survival, and apoptosis [37]. Finally, RELA and NFKB1 play an important role in melanoma progression, inflammation, and therapy resistance [38, 39]. These TFs represent potential biomarkers and therapeutic targets that could improve the understanding, diagnosis, and treatment of melanoma. Figure 3d presents the results of the transcription factor targets from enrichment analysis, identifying several target genes with known relevance to melanoma, including SOX10 target genes and GCCATNTTG YY1 Q6 (YY1). SOX10 is critical for melanocyte development and melanoma identity. It has been implicated in melanoma cell state transitions and phenotype switching [40]. YY1 contributes to human melanoma cell growth through modulating the p53 signaling pathway [41]. In Figure 3e, we retrieved various biological processes and pathways in melanoma progression. The top-enriched term is antigen processing and presentation of peptide antigen, particularly through MHC class I and II molecules. Multiple studies have shown that the antigen processing and presentation pathway is critical for melanoma immunogenicity and can influence both immune evasion and therapeutic response [42, 43]. Leukocyte differentiation is frequently identified as one of the top enriched processes in transcriptomic studies of melanoma, particularly in relation to immune response and tumor progression [44,45]. The Reactome pathway Signaling by Receptor Tyrosine Kinases contributes to melanoma progression and resistance to BRAF-targeted therapy through activation of EGFR, MET, PDGFR, and downstream MAPK and PI3K-AKT signaling [46]. We provide enrichment analysis results of Metascape based on DeepLIFT in Supplementary Metascape_Enrich1.

In Table 4, we report terms (i.e., drugs) and genes (i.e., drug targets) within IDG Drug Targets 2022. Vemurafenib and Dabrafenib are FDA-approved BRAF inhibitors that target the MAPK pathway in BRAF-mutant melanoma [47-49]. Sorafenib also targets RAF kinases, including ARAF, and has been investigated in melanoma clinical trials [50]. The method has also identified drugs such as Phenylalanine, a LAT1 (SLC7A5) inhibitor, and Metformin, a mitochondrial Complex I (NDUFA11) inhibitor, both associated with metabolic pathways implicated in melanoma [51, 52]. In Supplementary Table1_B, we list this enrichment analysis results for IDG Drug Targets 2022.

**Table 4:** Enriched terms from IDG Drug Targets 2022 via Enrichr were retrieved according to uploaded genes from DeepLIFT using A375 Cell Line, showing genes (column: genes) associated with drugs (column: term). The rank column shows the order of terms when retrieved.

| Rank | Term | Class | Genes | Approval Status / Study Phase | References |
| --- | --- | --- | --- | --- | --- |
| 5 | Vemurafenib | BRAF/MEK Pathway Inhibitor | ARAF | FDA Approved | [47] |
| 8 | Dabrafenib | BRAF/MEK Pathway Inhibitor | ARAF | FDA Approved | [48,49] |
| 15 | Sorafenib | VEGFR, BRAF, and c-KIT | ARAF | Clinical Trial | [50] |
| 1 | Phenylalanine | LAT1 inhibition | SLC7A5 | Preclinical Studies | [51] |
| 14 | Metformin | Mitochondrial Complex I inhibition | NDUFA11 | Clinical Trials | [52] |

#### 3.2.2. 451Lu Cell Line

Table 5 reports expressed genes within melanoma cell lines, obtained from the 60 human cancer cell lines (NCI-60) according to provided genes via all computational methods. Supplementary DataSheet2_A includes all genes generated by all computational methods, provided to Enrichr and Metascape. Among all evaluated methods, DeepLIFT and IG demonstrated superior performance compared to the others. Both methods identify a total of 31 expressed genes within the retrieved melanoma cell lines. In the MALME-3M cell line, both methods identified the same set of genes (DCT, S100A1, SLC16A6, APOD, CAPG, FXYD5, and RHOQ), with IG additionally detecting one more gene, L1CAM. In the MDA-MB-435 cell line, both methods identified a shared subset of genes (PLP1, CAPN3, and MAGEA3), with DeepLIFT uniquely detecting QDPR, COX7B, RPS6KA2, and TRMT12, while IG exclusively identified GPR143. In the SK-MEL-28 cell line, both methods identified a common set of genes (RCN3, SLC17A9, SLC16A6, LSAMP, APOD, CAPG, and RHOQ), with DeepLIFT uniquely detecting BFSP1, while IG additionally identified APP and DUSP10. In the SK-MEL-5 cell line, both methods identified a shared set of genes (DCT, S100A1, OR7E156P, SLC16A6, PLP1, CAPN3, and DHCR7), with DeepLIFT uniquely detecting SULT1C2 and TRMT12, while IG additionally identified GPR143, DUSP10, and L1CAM. Out of the 31 expressed genes identified across all cancer cell lines, 24 genes were shared by both DeepLIFT and IG, while DeepLIFT uniquely identified 7 genes (QDPR, COX7B, RPS6KA2, TRMT12, BFSP1, SULT1C2) and IG exclusively detected 4 genes (L1CAM, GPR143, APP, DUSP10). The worst-performing methods were LIMMA with 24 expressed genes within MALME-3M, MDA-MB435, SKMEL28, and SKMEL5 melanoma cell lines. Supplementary Table2_A includes enrichment analysis results for NCI-60 Cancer Cell Lines, obtained from Enrichr.

**Table 5:**
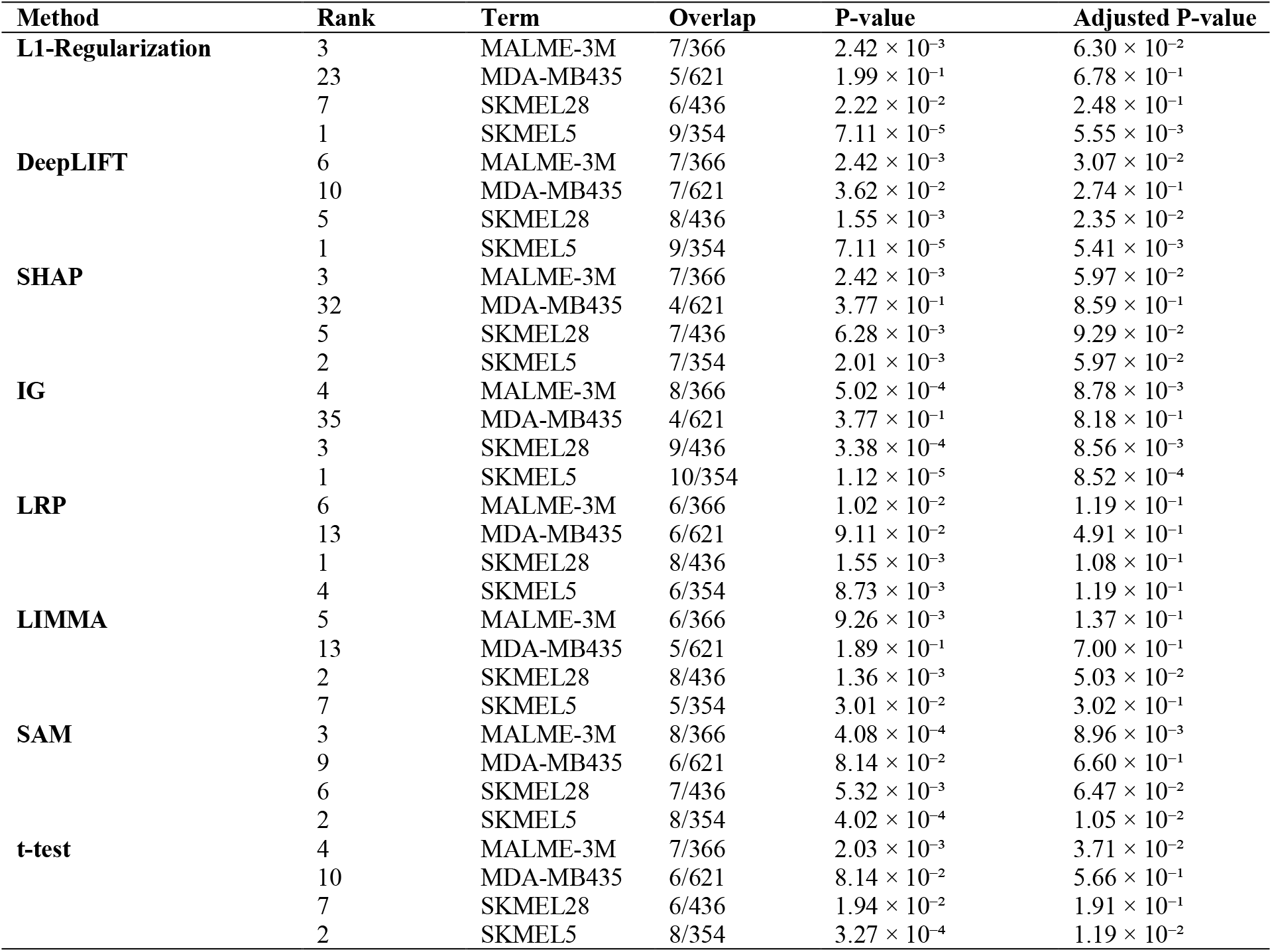
NCI-60 Cancer Cell Lines via Enrichr according to the genes produced by each method when using 451Lu Cell Line.

Figure 4a displays an UpSet plot illustrating the intersection of gene sets identified by all computational methods. Methods integrated within our approach identified a notable number of unique genes, ranging from 19 to 13, with the exception of L1-Regularization including only two unique genes. In contrast, traditional bioinformatic approaches identified fewer unique genes— only one for the t-test, none for SAM, and 10 for LIMMA. This points out to a key advantage of our DL-based approach, which not only recovers established biological signals but also identifies additional, potentially novel genes often missed by conventional approaches. In Supplementary DataSheet2_B, we included the UpSet plot of gene lists provided by DeepLIFT and IG with bioinformatics tools for 451Lu Cell Lines.

**Figure 4:**
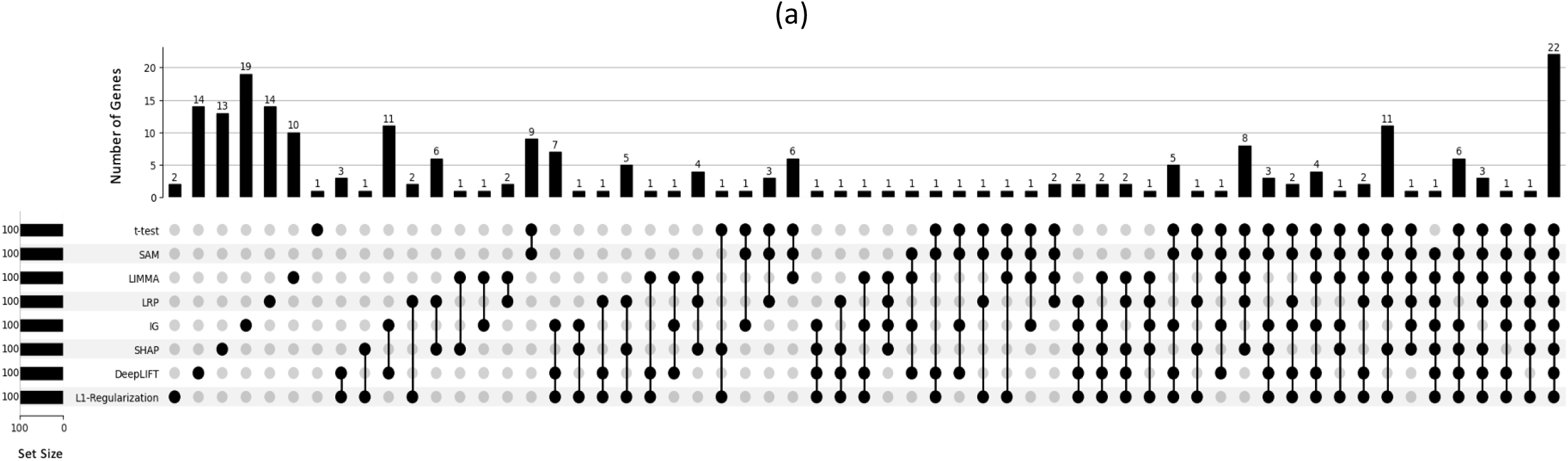
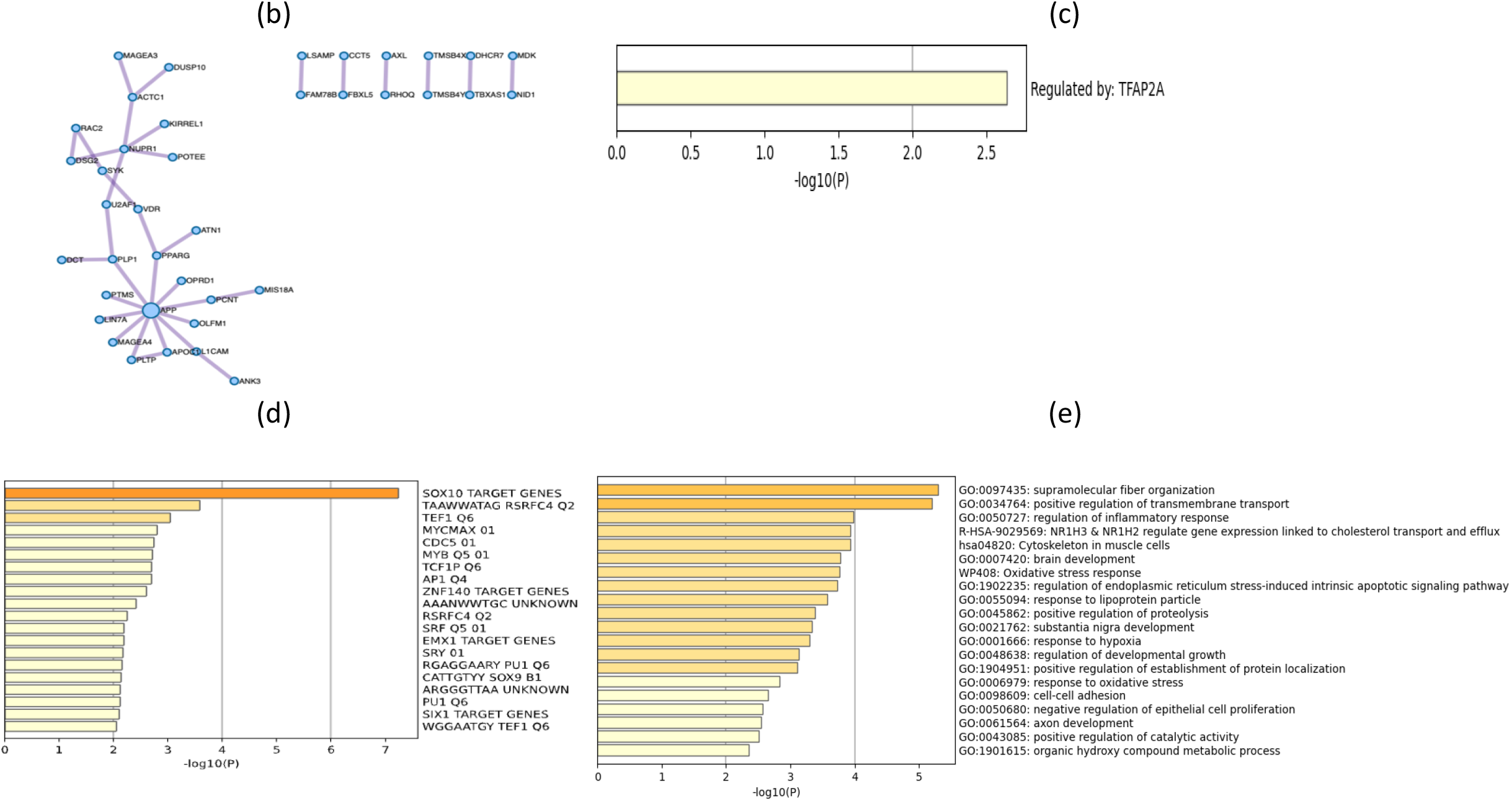
(a) UpSet plot of gene lists provided by all the computational methods when using 451Lu Cell Line. (b) Two clusters identified in PPI network pertaining to 451Lu Cell Line. (c) A retrieved transcription factor based on genes from DeepLIFT when provided to Metascape. (d) Enrichment analysis pertaining to twenty transcription factor targets for studied 451Lu Cell Line. (e) Process and pathway enrichment analysis provided by Metascape according to the 100 genes of DeepLIFT using 451Lu Cell Line.

While both DeepLIFT and IG performed well, the IG-unique gene set demonstrated a stronger association with melanoma biology, as evidenced by the presence of L1CAM, GPR143, and DUSP10, all these genes are with known or suspected roles in melanoma. Therefore, to unveil biological insights within melanoma drug responses, we focused on the IG-derived gene set provided to Metascape enrichment analysis. Figure 4b displays The following 39 genes obtained from protein–protein interaction (PPI): POTEE, FAM78B, KIRREL1, MIS18A, NUPR1, FBXL5, RHOQ, CCT5, DUSP10, OLFM1, TMSB4Y, LIN7A, VDR, U2AF1, TMSB4X, TBXAS1, SYK, RAC2, PTMS, PPARG, PLTP, PLP1, PCNT, OPRD1, NID1, MDK, MAGEA4, MAGEA3, LSAMP, L1CAM, DSG2, ATN1, DHCR7, DCT, AXL, APP, APOC1, ANK3, ACTC1.

Figure 4c reports 1 TF: TFAP2A, as it plays a role as a transcriptional coactivator in melanoma, particularly by regulating immune-related genes (like MHC II) and influencing tumor immunogenicity [53]. Figure 4d presents the enrichment analysis results for transcription factor targets, identifying several targets with known relevance to melanoma, including SOX10 target genes and MYCMAX 01. As mentioned, SOX10 is essential for melanocyte development and maintaining melanoma identity. It controls important melanocytic genes like MITF, TYR, and DCT and plays a role in melanoma cell state changes and phenotype switching [40]. The c-Myc/MAX complex promotes melanoma growth and spread by binding to the E-box motif (CACGTG) and activating genes that drive cell proliferation. This binding is also common in drug-tolerant melanoma cells that resist BRAF/MEK inhibitors, highlighting its role in therapy resistance [54]. Genes provided via IG are related to various biological processes and pathways in melanoma progression, as illustrated in Figure 4e. The analysis reports that the regulation of inflammatory response is the highest biological processes related to melanoma. Disruption or abnormal regulation of supramolecular fiber organization has been associated with increased melanoma aggressiveness, enhanced metastatic potential, and resistance to therapy [55]. In addition, Regulation of inflammatory response is also reported, and it is relevant and actively studied in the context of melanoma development, progression, and treatment, especially due to its implications in immunotherapy [56]. We include enrichment analysis results of Metascape based on DeepLIFT in Supplementary Metascape_Enrich2.

In Table 6, we present drugs and their associated expressed genes identified within the IDG Drug Targets 2022 database, focusing on their relevance in melanoma treatment and resistance. Nelfinavir, an HIV protease inhibitor, has demonstrated preclinical and early clinical activity against melanoma and is being investigated for use in drug-resistant cases [57]. Imatinib, which targets c-KIT mutations commonly found in specific melanoma subtypes such as AM, is FDA-approved for treating c-KIT mutant melanoma [58,59]. In addition, the analysis identified several multi-kinase inhibitors, including Cabozantinib, Crizotinib, Gefitinib, Axitinib, and Dasatinib, which target receptor tyrosine kinases (RTKs) such as AXL, VEGFRs, MET, and Src kinases. These RTKs play critical roles in melanoma progression and therapy resistance, highlighting the potential of these drugs for therapeutic intervention in advanced or resistant melanoma cases [60–64]. In Supplementary Table2_B, we list this enrichment analysis results for IDG Drug Targets 2022.

**Table 6:** Enriched terms from IDG Drug Targets 2022 via Enrichr were retrieved according to uploaded genes from DeepLIFT using 451Lu Cell Line, showing genes (column: genes) associated with drugs (column: term). The rank column shows the order of terms when retrieved.

| Rank | Term | Class | Genes | Approval Status / Study Phase | References |
| --- | --- | --- | --- | --- | --- |
| 11 | Nelfinavir | ER stress / PI3K-Akt inhibition | OPRD1; TBXAS1 | Clinical trials | [57] |
| 54 | Imatinib | KIT (c-KIT) | SYK; TBXAS1 | Clinical trials | [58,59] |
| 69 | Cabozantinib | AXL, MET, VEGFR2 | AXL | Clinical trials | [60] |
| 100 | Crizotinib | ALK, MET, AXL, RON | SYK; AXL | Clinical trials | [61] |
| 112 | Gefitinib | EGFR | AXL | Clinical trials | [62] |
| 114 | Axitinib | VEGFR1-3 | AXL | Clinical trials | [63] |
| 120 | Dasatinib | BCR-ABL / Src (SYK modulator) | SYK | Clinical trials | [64] |

### 3.3. Classification Results

In this section, we assess the performance from a classification perspective, comparing those gene sets derived from methods our DL-based approach with baseline methods. Each gene set was used in the gene expression dataset to train three machine learning models: support vector machines (SVM), random forest (RF), and a neural network (NN). We utilized five-fold cross-validation to report predictive performance incorporating F1 and balanced accuracy (BAC) performance metrics.

When considering GSE108383 dataset with A375 Cell Line, Figure 5a demonstrates that the IG method in our approach coupled with SVM and NN obtained the highest BAC of 0.980 (tie with our method LRP) and 1.00, respectively, while t-test coupled with RF obtained the highest BAC of 1.00. In terms of the BAC performance measure, the overall loss for models when utilizing our best method IG is (1-0.98) + (1-0.98) + (1-1) = 4%. The second-best method is DeepLIFT in which the total loss for employed models is (1-0.974) + (1-0.99) + (1-0.99) = 4.6%. The overall loss for models when utilizing the best bioinformatics-based tool (LIMMA) is (1-0.97) + (1-0.99) + (1-0.98) = 6%. The worst performance result is related to t-test in which the total loss is (1-0.943) + (1-1) + (1-0.971) = 8.6%. These results reveal 2% and 1.4% BAC performance improvements of IG and DeepLIFT, respectively, over the existing tool, LIMMA. For the F1 performance metric (see Figure 5b), our IG method when utilized in all three models contributed to (1-0.979) + (1-0.979) + (1-1) = 4.2% total loss. The second-best method is DeepLIFT in our approach in which the total loss is (1-0.968) + (1-0.99) + (1-0.99) = 5.2%. LIMMA, the best among existing tools, had a total loss of (1-0.967) + (1-0.99) + (1-0.979) = 6.4%. These results demonstrate 2.2% and 1.2% performance improvements over LIMMA when F1 is considered. Tables S1 and S2 in Supplementary Additional File report these performance results in tabular formats.

**Figure 5:**
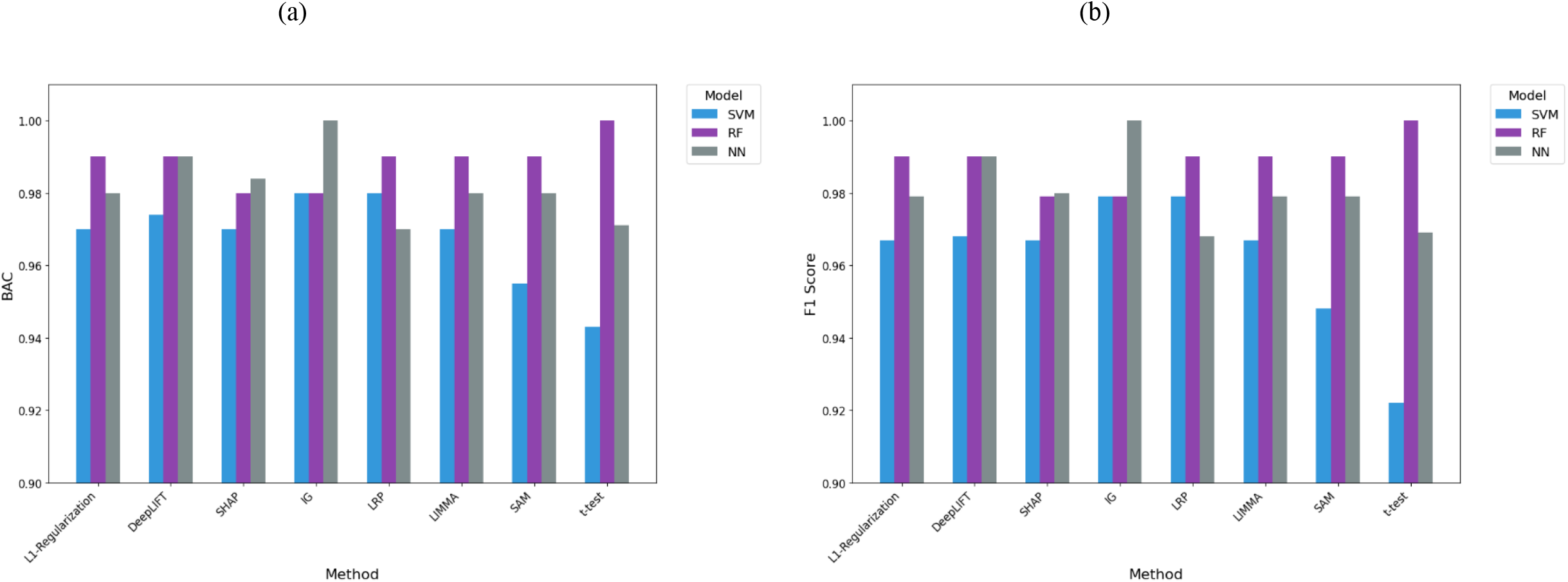
(a) BAC of each model during five-fold cross-validation when coupled with gene sets of each method according to A375 Cell Line. (b) F1 results of each model during five-fold cross-validation when coupled with gene sets produced via each method when using A375 Cell Line.

For the 451Lu cell line dataset when utilizing the BAC performance measure (see Figure 6a), the gene set produced by IG method in our approach achieved the highest BAC of 1.00 when employing RF and NN. The second-best method is t-test yielding the highest BAC of 0.976 (tie with SAM) when coupled with SVM while having a tie with IG when RF is utilized. Specifically, IG method in our approach when coupled with all models had a total loss of (1-0.97) + (1-1) + (1-1) = 3%. The second-best method is related to t-test in the baseline approach, resulting in a total loss of (1-0.976) + (1-1) + (1-0.991) = 3.3%. These results demonstrate a BAC performance improvement of 0.3% over the existing approach utilizing t-test method. The worst-performing method is related to SHAP when used in our approach in which producing a total loss of (1-0.97) + (1-0.982) + (1-0.984) = 6.4%. For the F1 performance metric in Figure 6b, IG method in our approach had the highest F1 of 1 when RF and NN are utilized. On the other hand, t-test had the highest F1 of 0.983 and 1 when coupled with SVM (tie with SAM) and RF (tie with IG in our approach), respectively. It can be seen from Figure 6b that IG in our approach had the lowest loss of (1-0.979) + (1-1) + (1-1) = 2.1%, while t-test had a higher loss of (1-0.983) + (1-1) + (1-0.991) = 2.6%. In other words, we have a 0.5% F1 performance improvement over the existing baseline method t-test. These findings highlight the advantage of our approach with IG over both other DL-based and traditional bioinformatics methods. Tables S3 and S4 in Supplementary Additional File include these performance results in a tabular format pertaining to 451Lu Cell Line.

**Figure 6.**
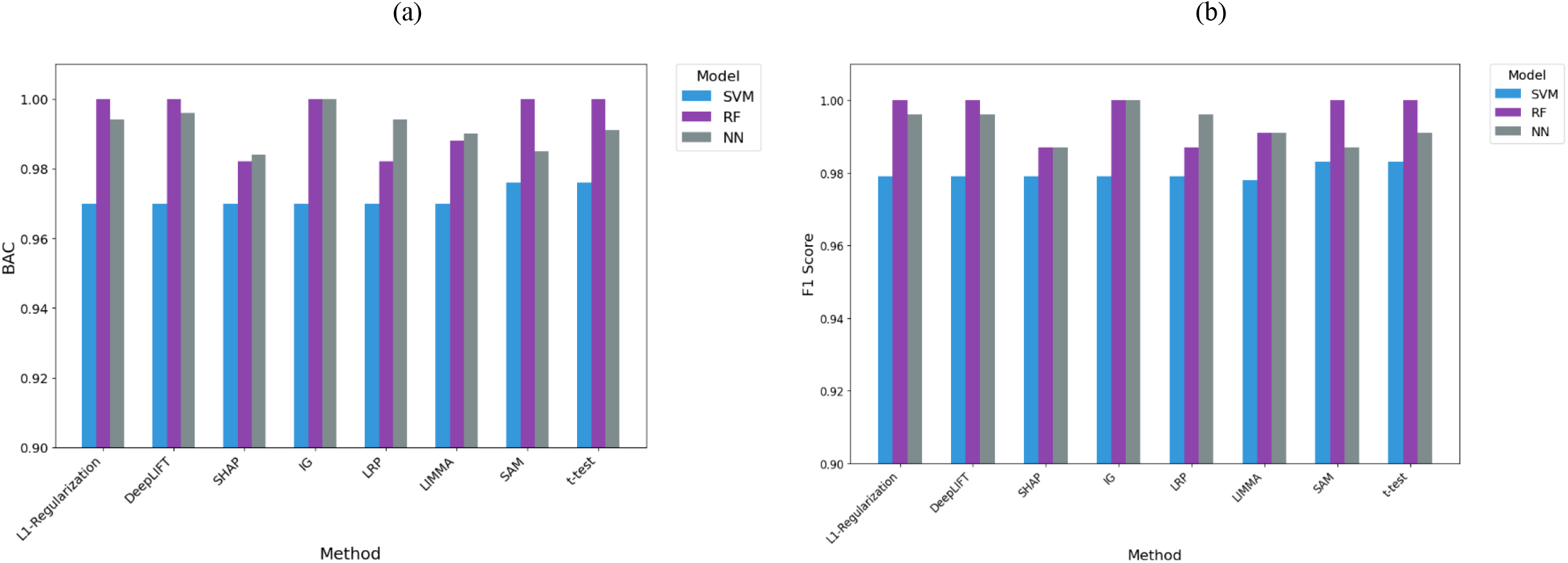
(a) The BAC of each model during five-fold cross-validation by the gene set of each method using 451Lu cell line (b) F1 results of each model during five-fold cross-validation when coupled with gene sets produced via each method when using 451Lu cell line.

## 4. Discussion

To understand the mechanisms underlying drug resistance in melanoma and thereby improve the treatment outcomes and survival rates, we present a DL-based approach that works as follows. First, we downloaded scRNA-seq gene expression data from GEO repository pertaining to two melanoma cell lines, each provided to a fully connected neural network with five adapted methods (L1-Regularization, DeepLIFT, SHAP, IG and LRP)) to discriminate between BRAFi-resistant and parental cell lines. Then, we select top-100 genes with highest weights as described in Section 2.2.2. We provide top-100 genes derived from our approach as well as baseline methods for enrichment analysis. Results demonstrated that methods in our approach identified more expressed genes in four well-established melanoma cell lines, followed by identifying (1) FDA approved drugs in melanoma; and (2) drug targets, transcription factors, biological processes and pathways. Results from a classification perspective demonstrated that gene sets derived from adapted methods in our approach when coupled with machine learning algorithms improved the prediction performance and these results indicate that produced genes are discriminative.

It can be seen from results in Section 3.2. that DeepLIFT and IG, the best-performing methods in our approach, identified genes such as SOX10, ARAF, SLC7A5, DCT, and AXL, which are known to play critical roles in melanoma progression, phenotype switching, and drug resistance. SOX10 has been implicated in melanoma cell state transitions and therapy resistance. Similarly, ARAF and AXL are associated with resistance to BRAF/MEK inhibitors, while SLC7A5 (LAT1) has been linked to metabolic reprogramming in melanoma. In addition to identifying critical genes, our enrichment analysis revealed key pathways such as antigen processing and presentation, leukocyte differentiation, and signaling by receptor tyrosine kinases (RTKs). These pathways are well-documented in melanoma research, particularly in the context of immune evasion, tumor progression, and therapy resistance. For example, the antigen processing and presentation pathway is crucial for melanoma immunogenicity and influences therapeutic responses to inhibitors. The identification of these pathways demonstrates the biological relevance and robustness of our approach.

Our DL-based approach demonstrated clear advantages over traditional bioinformatics tools such as LIMMA, SAM and t-test. These traditional methods rely on linear assumptions and limited capacity to capture complex interactions within high-dimensional data. In contrast, adapted methods in our DL-based approach captures non-linear relationships, enabling the identification of a broader set of genes with strong biological relevance. For instance, in the A375 cell line dataset, IG identified 32 expressed genes in the four well-established melanoma cell lines while LIMMA identified 29 expressed genes. In terms of 451Lu cell line dataset, IG in our approach identified 31 expressed genes in the four well-established cell lines while t-test identified 27 expressed genes. Moreover, our analysis highlighted several FDA-approved drugs and potential therapeutic candidates for melanoma. Notably, Vemurafenib and Dabrafenib, both BRAF inhibitors, were identified as key drugs associated with genes such as ARAF. Additionally, Sorafenib, which targets RAF kinases, and Nelfinavir, an HIV protease inhibitor with emerging anti-melanoma properties, were identified as potential therapeutic options. These findings align with current clinical practices and highlight the potential of our DL-based approach to aid in drug repurposing for melanoma.

Moreover, transcription factors such as MITF, RELA, E2F1, and TFAP2A were identified as critical regulators of melanoma progression and drug resistance. These TFs not only serve as biomarkers but also represent potential targets for therapeutic intervention. For instance, MITF regulates melanocyte lineage survival, while E2F1 is involved in melanoma cell proliferation and apoptosis. RELA and NFKB1 are central to inflammatory signaling and therapy resistance, providing avenues for targeted therapies.

From a classification perspective, gene sets selected by IG within our approach coupled with machine learning algorithms demonstrated superior predictive performance compared to the best-performing existing methods. In the A375 cell line dataset, IG had a total loss of 4.2% and 4%, measured using F1 and BAC, respectively, while LIMMA had a total loss of 6.4% and 6% according to F1 and BAC, respectively. For the 451Lu cell line dataset, IG resulted in a total loss of 2.1% and 3% measured using F1 and BAC, respectively, while t-test had a total loss of 2.6% and 3.3% based on F1 and BAC, respectively.

## 5. Conclusions and Future Work

This study presents a novel DL-based computational approach to uncover critical genes, pathways, drugs and therapeutic targets associated with drug resistance in melanoma. Compared to existing tools, our approach effectively identifies biologically and clinically relevant genes, as demonstrated by its superiority from biological and classification perspectives. Biological assessment using two melanoma cell lines datasets from the GEO repository demonstrate that methods in our approach identifying more expressed genes in four widely studied melanoma cell lines, including MALME-3M, MDA-MB435, SK-MEL-28, and SK-MEL-5. Biological Key findings include the identification of critical genes such as SOX10, ARAF, SLC7A5, and AXL, as well as transcription factors like MITF and RELA, which are linked to melanoma progression and drug resistance. Our DL-based approach also identified FDA-approved drugs, including Vemurafenib and Dabrafenib, as well as potential therapeutic candidates like Sorafenib and Nelfinavir, demonstrating its potential in speeding up the drug discovery process. Assessment from a classification point of view demonstrates that adapting integrated gradients (IG) in our approach resulted in 2.2% and 0.5% overall performance improvements when compared to the best-performing baselines based on A375 and 451Lu cell line datasets. These results demonstrate the capability of our DL-based approach in (1) reducing the search space pertaining to drugs, therapeutic targets, biomarker genes, biological processes and pathways in melanoma; and (2) identifying discriminant gene sets that can guide in predicting melanoma drug response.

Future work includes (1) employing our DL-based approach to unveil shared molecular mechanisms pertaining to melanoma and other cancer types; (2) identifying gene signatures for melanoma patients treated with multiple targeted inhibitors (3) incorporating agentic AI with our DL-based approach to improve the outcomes of different cancer types from both biological and classification perspectives.

## Author Contributions

S.A.: methodology, software, visualization, investigation, writing—original draft preparation. T.T.: conceptualization, methodology, data curation, supervision, writing—reviewing and editing. Y.-h.T.: validation, writing—reviewing and editing. All authors have read and agreed to the submitted version of the manuscript.

## Funding

This study received no funding.

## Data Availability Statement

Data are contained within the article.

## Conflict of Interest

The authors declare no conflict of interest.

## Notes

### Competing Interest Statement

The authors have declared no competing interest.

