## Supplementary material for "De novo single-cell biological analysis of drug resistance in human melanoma through a novel deep learning-powered approach": Additional File.docx

**Table S1:** Five-fold cross-validation results on GSE108383 dataset with A375 Cell Line using gene sets by each method, provided to three machine learning methods. BAC is balanced accuracy.

| **Method** | **Model** | **BAC** | **F1** |
| --- | --- | --- | --- |
| L1-Regularization | SVM | 0.970 ± 0.040 | 0.967 ± 0.044 |
|  | RF | 0.990 ± 0.020 | 0.990 ± 0.021 |
|  | NN | 0.980 ± 0.025 | 0.979 ± 0.026 |
| DeepLIFT | SVM | 0.974 ± 0.039 | 0.968 ± 0.044 |
|  | RF | 0.990 ± 0.020 | 0.990 ± 0.021 |
|  | NN | 0.990 ± 0.020 | 0.990 ± 0.021 |
| Shap | SVM | 0.970 ± 0.040 | 0.967 ± 0.044 |
|  | RF | 0.980 ± 0.025 | 0.979 ± 0.026 |
|  | NN | 0.984 ± 0.021 | 0.980 ± 0.025 |
| IG | SVM | 0.980 ± 0.025 | 0.979 ± 0.026 |
|  | RF | 0.980 ± 0.025 | 0.979 ± 0.026 |
|  | NN | 1.000 ± 0.000 | 1.000 ± 0.000 |
| LRP | SVM | 0.980 ± 0.025 | 0.979 ± 0.026 |
|  | RF | 0.990 ± 0.020 | 0.990 ± 0.021 |
|  | NN | 0.970 ± 0.025 | 0.968 ± 0.026 |
| LIMMA | SVM | 0.970 ± 0.040 | 0.967 ± 0.044 |
|  | RF | 0.990 ± 0.020 | 0.990 ± 0.021 |
|  | NN | 0.980 ± 0.025 | 0.979 ± 0.026 |
| SAM | SVM | 0.955 ± 0.032 | 0.948 ± 0.035 |
|  | RF | 0.990 ± 0.020 | 0.990 ± 0.021 |
|  | NN | 0.980 ± 0.025 | 0.979 ± 0.026 |
| t-test | SVM | 0.943 ± 0.037 | 0.922 ± 0.050 |
|  | RF | 1.000 ± 0.000 | 1.000 ± 0.000 |
|  | NN | 0.971 ± 0.024 | 0.969 ± 0.025 |

**Table S2:** Balanced accuracy (BAC) and F1 performance difference **∆** for each method across the three models from Table S1. The higher **∆** the results the worse the performance is. The best results are shown in bold.

| **Method** | **∆BAC** | **∆F1** |
| --- | --- | --- |
| L1-Regularization | 6.0% | 6.4% |
| DeepLIFT | 4.6% | 5.2% |
| SHAP | 6.6% | 7.4% |
| IG | **4.0%** | **4.2%** |
| LRP | 6.0% | 6.3% |
| LIMMA | 6.0% | 6.4% |
| SAM | 7.5% | 8.3% |
| t-test | 8.6% | 10.9% |

**Table S3:** Five-fold cross-validation results on GSE108383 dataset with 451Lu Cell Line using gene sets by each method, provided to three machine learning methods. BAC is balanced accuracy.

| **Method** | **Model** | **BAC** | **F1** |
| --- | --- | --- | --- |
| L1-Regularization | SVM | 0.970 ± 0.019 | 0.979 ± 0.013 |
|  | RF | 1.000 ± 0.000 | 1.000 ± 0.000 |
|  | NN | 0.994 ± 0.012 | 0.996 ± 0.009 |
| DeepLIFT | SVM | 0.970 ± 0.019 | 0.979 ± 0.013 |
|  | RF | 1.000 ± 0.000 | 1.000 ± 0.000 |
|  | NN | 0.996 ± 0.009 | 0.996 ± 0.009 |
| SHAP | SVM | 0.970 ± 0.019 | 0.979 ± 0.013 |
|  | RF | 0.982 ± 0.024 | 0.987 ± 0.017 |
|  | NN | 0.984 ± 0.014 | 0.987 ± 0.011 |
| IG | SVM | 0.970 ± 0.019 | 0.979 ± 0.013 |
|  | RF | 1.000 ± 0.000 | 1.000 ± 0.000 |
|  | NN | 1.000 ± 0.000 | 1.000 ± 0.000 |
| LRP | SVM | 0.970 ± 0.019 | 0.979 ± 0.013 |
|  | RF | 0.982 ± 0.024 | 0.987 ± 0.017 |
|  | NN | 0.994 ± 0.012 | 0.996 ± 0.009 |
| LIMMA | SVM | 0.970 ± 0.019 | 0.978 ± 0.013 |
|  | RF | 0.988 ± 0.014 | 0.991 ± 0.011 |
|  | NN | 0.990 ± 0.012 | 0.991 ± 0.010 |
| SAM | SVM | 0.976 ± 0.012 | 0.983 ± 0.009 |
|  | RF | 1.000 ± 0.000 | 1.000 ± 0.000 |
|  | NN | 0.985 ± 0.012 | 0.987 ± 0.011 |
| t-test | SVM | 0.976 ± 0.012 | 0.983 ± 0.009 |
|  | RF | 1.000 ± 0.000 | 1.000 ± 0.000 |
|  | NN | 0.991 ± 0.011 | 0.991 ± 0.011 |

**Table S4:** Balanced accuracy (BAC) and F1 performance difference **∆** for each method across the three models from Table S3. The higher **∆** the results the worse the performance is. The best results are shown in bold.

| **Method** | **∆BAC** | **∆F1** |
| --- | --- | --- |
| L1-Regularization | 3.6% | 2.5% |
| DeepLIFT | 3.4% | 2.5% |
| SHAP | 6.4% | 4.7% |
| IG | **3.0%** | **2.1%** |
| LRP | 5.4% | 3.8% |
| LIMMA | 5.2% | 4.0% |
| SAM | 3.9% | 3.0% |
| t-test | 3.3% | 2.6% |
